# The metabolic reprogramming of T cells controls airway remodeling in severe asthma

**DOI:** 10.64898/2026.03.19.712985

**Authors:** Hope Steele, Elizabeth Kato, Garrison Dell, Maeve Fink, Andrew Ghastine, Ashley Willicut, Hilde Cheroutre, Mitchell Kronenberg, Rana Herro

## Abstract

Mixed granulocytic asthma (MGA) is a severe Th2-low endotype, characterized by high Th17/neutrophilic burden and exacerbated airway remodeling. Both features confer resistance to inhaled corticosteroids, and typical asthma treatments. Thus, MGA is an enormous public health burden. Gaps in knowledge include how Th17 cells induce pathological tissue remodeling, and how Th17 differentiation occurs in response to allergens. We generated a Th2-low murine model of asthma that recapitulates major features of human MGA namely, heightened airway reactivity to methacholine, Th17/neutrophilic inflammation, airway remodeling, and resistance to corticosteroid treatment. Two specific biomarkers enriched in human MGA, the TNF superfamily member 14 (aka LIGHT), and the mitochondrial oxidative phosphorylation (OXPHOS) pathway, are upregulated in this model. We show OXPHOS promotes the metabolic reprograming of Th17 cells, to produce LIGHT that controls airway remodeling. Mechanistically, OXPHOS regulates RORγt expression and the subsequent transcriptional network to program survival and differentiation of Th17 cells, whereas LIGHT drives airway remodeling by activating the MMP9-dependent TGFβ pathway. Additionally, OXPHOS^+^Th17 cells promote the expression of osteopontin necessary for fibroblast activation. LIGHT antagonistic blockade reduces airway remodeling, whereas OXPHOS chemical inhibition reduces Th17 cells and neutrophilia. Importantly, the dual blockade of LIGHT and OXPHOS reverses all features of MGA and reciprocally increase the numbers of Treg cells. Thus, the dual blockade of LIGHT and OXPHOS constitutes a promising target for clinical interventions in human MGA, possibly extending to other Th17-driven fibrotic diseases.

## INTRODUCTION

Despite decades of research, severe asthma remains strongly associated with substantial morbidities and mortalities (1). Non-Th2 mixed granulocytic asthma (MGA) is a severe endotype resistant to corticosteroid and long-acting β-agonist treatments. MGA is due to enhanced airway remodeling (2–18) and increased Th17/neutrophilic inflammation (19–27). Understanding critical signaling pathways driving airway remodeling and Th17 cells will help define novel targets for successful clinical intervention; however, how these immunological shifts occur remains poorly understood. Steroids kill Th2 cells (28) and eosinophils (29), but neutrophils and Th17 cells are resistant (26). Additionally, airway remodeling requires anti-fibrotic drugs for treatment rather than anti-inflammatory therapies or bronchodilators. Clinical trials targeting Th2 cytokines failed to treat severe asthma, arguably due to stronger association with Th17/neutrophilic inflammation than Th2 cells (30–40).

Emerging data show that TNF superfamily member 14 (TNFSF14, aka LIGHT) and the mitochondrial oxidative phosphorylation (OXPHOS) pathway correlate with non-Th2 asthma in humans (41, 42), and are increased in asthmatic-patients with high Th17 cells in their sputum (42, 43). We therefore hypothesized that LIGHT promotes airway remodeling while OXPHOS promotes Th17 inflammation, synergizing together to drive MGA. Our group pioneered work showing LIGHT is a central regulator of fibrosis and can directly stimulate normal human lung epithelial cells and fibroblasts to induce remodeling (44–47). LIGHT signaling through its receptor LTβR is essential for fibroproliferation, smooth muscle cell non-canonical activation, and epithelial cell hyperplasia (45, 46, 48). LIGHT inhibition protected from pathological airway remodeling associated with idiopathic pulmonary fibrosis (46, 49). In a mouse model of eosinophilic asthma (EA), abrogating LIGHT signaling decreased TGFβ production by macrophages, IL-13 expression by eosinophils, and subsequent airway inflammation (49). Furthermore, LIGHT can program a steroid-resistant signature in human lung epithelial cells (50).

Here, we show LIGHT controls airway remodeling in MGA. We used *Il4rα^-/-^* mice to impose defective IL4/IL13 signaling and failure to mount a Th2 response, to mimic human MGA. We demonstrate upon chronic exposure to house dust mite (HDM), *Il4rα^-/-^* exhibited increased Th17/neutrophilic lung inflammation, exacerbated airway remodeling and hyperreactivity to methacholine, and resistance to corticosteroid and anti-IL17 therapies, recapitulating seminal features of MGA. Prior literature suggests that OXPHOS controls Th17 cell anti-apoptotic survival (51) and IL-17 production (52). In asthmatic patients, OXPHOS is upregulated, coinciding with high Th17 cellularity in the sputum (42, 43). One key gap in knowledge is how OXPHOS regulates Th17 cells in MGA. We provide a mechanistic connection between OXPHOS, increased Th17 cells, and lung fibrosis. Our study validates and integrates LIGHT and OXPHOS in MGA pathogenesis. Furthermore, we show LIGHT/OXPHOS signature is induced in several human Th17-driven fibrotic diseases, including multiple sclerosis, rheumatoid arthritis, chronic obstructive pulmonary disease, psoriasis, psoriatic arthritis, systemic lupus, idiopathic pulmonary fibrosis, in addition to severe asthma.

## RESULTS

### IL4Rα deficiency generates a non-Th2 model of mixed granulocytic asthma (MGA)

Recent literature shows upregulation of LIGHT and OXPHOS in human asthmatics with high Th17 cellularity (41, 42). We focused on generating a mouse model that recapitulates seminal features of human MGA, including increased Th17 cellularity, neutrophilia, enhanced airway remodeling and hyperreactivity, and resistance to steroids and anti-IL17 therapies. Because severe asthmatic patients are resistant to corticosteroids, which downregulate IL4Rα signaling (53), we analyzed mice with ablated IL4Rα expression to impose defective IL4/IL13 signaling and disable Th2 responses. After chronic exposure to HDM over three consecutive weeks, we show *Il4rα^-/-^* lungs display features of severe MGA, including enhanced airway remodeling as evidenced by sub-epithelial fibrosis and collagen deposition, and inflammation (Fig. 1A), in addition to an increase in IL-17 levels and Th17 cellularity (Fig. 1C-D). Compared to wildtype mice that generate a Th2 EA model, HDM-induced *Il4rα^-/-^* mice exhibit significantly higher airway hyperreactivity in response to methacholine (Fig. 1B), and enhanced airway neutrophilia with comparable eosinophilia (Fig. 1D) and IL-13 levels (although IL-13 signaling is defective due to the absence of its receptor) (Fig. 1C). Treatment with dexamethasone did not reduce airway remodeling or neutrophilic inflammation (Fig. 1E-F). Likewise, IL-17 blockade did not reduce MGA pathology (Fig. 1G). Collectively, these features recapitulate the critical characteristics of human non-Th2 MGA (54).

**Figure 1:**
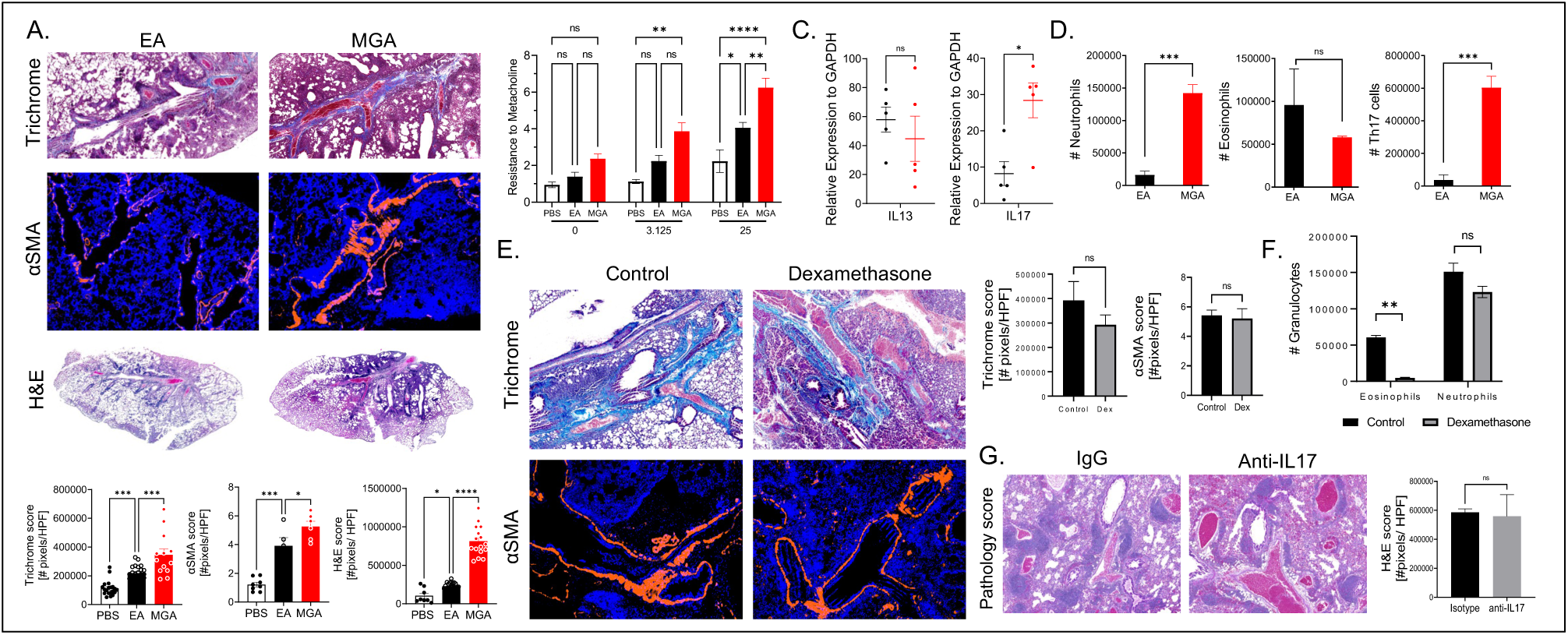
IL4Rα deficiency generates a new model of mixed granulocytic asthma (MGA). To model EA or MGA respectively, Balb/c or *Il4ra^-/-^* mice were given five intranasal (i.n.) challenges with 100ug house dust mite (HDM) extract every week (followed by two days resting) for three consecutive weeks. A final i.n. HDM challenge was given 24 hours prior to harvest. (**A**) Lung sections were stained and representative Masson trichrome (collagen), α-smooth muscle actin (αSMA), and hematoxylin and eosin (H&E) sections and accompanying quantifications are shown. (**B**) Airway resistance to increasing doses of methacholine exposure (6-8 mice per group). **(C)** Expression of mRNA for IL13 and IL17, assessed by quantitative PCR analyses of lung samples. mRNA expression was calculated relative to GAPDH for 5 mice per group. (**D**) Number of neutrophils or eosinophils in the left lung lobe of EA or MGA induced mice, as assessed by flow cytometry. (**E**) MGA-induced animals were treated with 50ug dexamethasone intratracheally one hour prior to each allergen challenge, beginning on day 7. Lung sections were stained and representative Masson trichrome and αSMA sections and accompanying quantifications are shown. (**F**) Number of neutrophils or eosinophils in the left lung lobe of MGA induced mice treated with or without dexamethasone, as assessed by flow cytometry. (**G**) MGA-induced animals were treated with 100ug anti-IL17A or matching isotype control i.p. beginning on day 7 and continuing every other day until harvest. Lung sections were stained, and representative H&E sections and accompanying quantifications are shown. Trichrome data show area of interest values covering full lung section of 6 individual mice per condition. αSMA data show mean values of all bronchi scored from 6 mice per condition. Data are representative of 3 independent experiments. *P < 0.05, **P < 0.005, ***P < 0.0005, ****P < 0.00005; HPF, high-power field; ns, nonsignificant.

### Th17 cells are essential drivers of MGA pathology

Using anti-CD4/CD8 depleting antibodies during disease initiation and throughout, we show *Il4rα^-/-^* mice lacking T cells are protected from HDM-induced MGA (Fig. 2). Airway remodeling drastically decreases in the absence of T cells (Fig. 2A), in addition to a decrease in neutrophilia (Fig. 2B). Equal numbers of Th2 (CD45.2^+^CD3^+^CCR6^-^CCR4^+^CXCR3^-^CCR10^-^) or Th17 cells (CD45.2^+^CD3^+^CCR6^+^CCR4^+^CXCR3^-^CCR10^-^) were sorted from HDM challenged BALB/c or *Il4ra^-/-^* animals and transferred intratracheally into naïve *Rag1^-/-^* hosts. All recipient mice were challenged intranasally with HDM. Th17-recipients show enhanced airway remodeling compared to mice receiving equivalent numbers of Th2 cells (Fig. 2C), emphasizing increased lung fibrosis with Th17 compared to Th2 cells.

**Figure 2:**
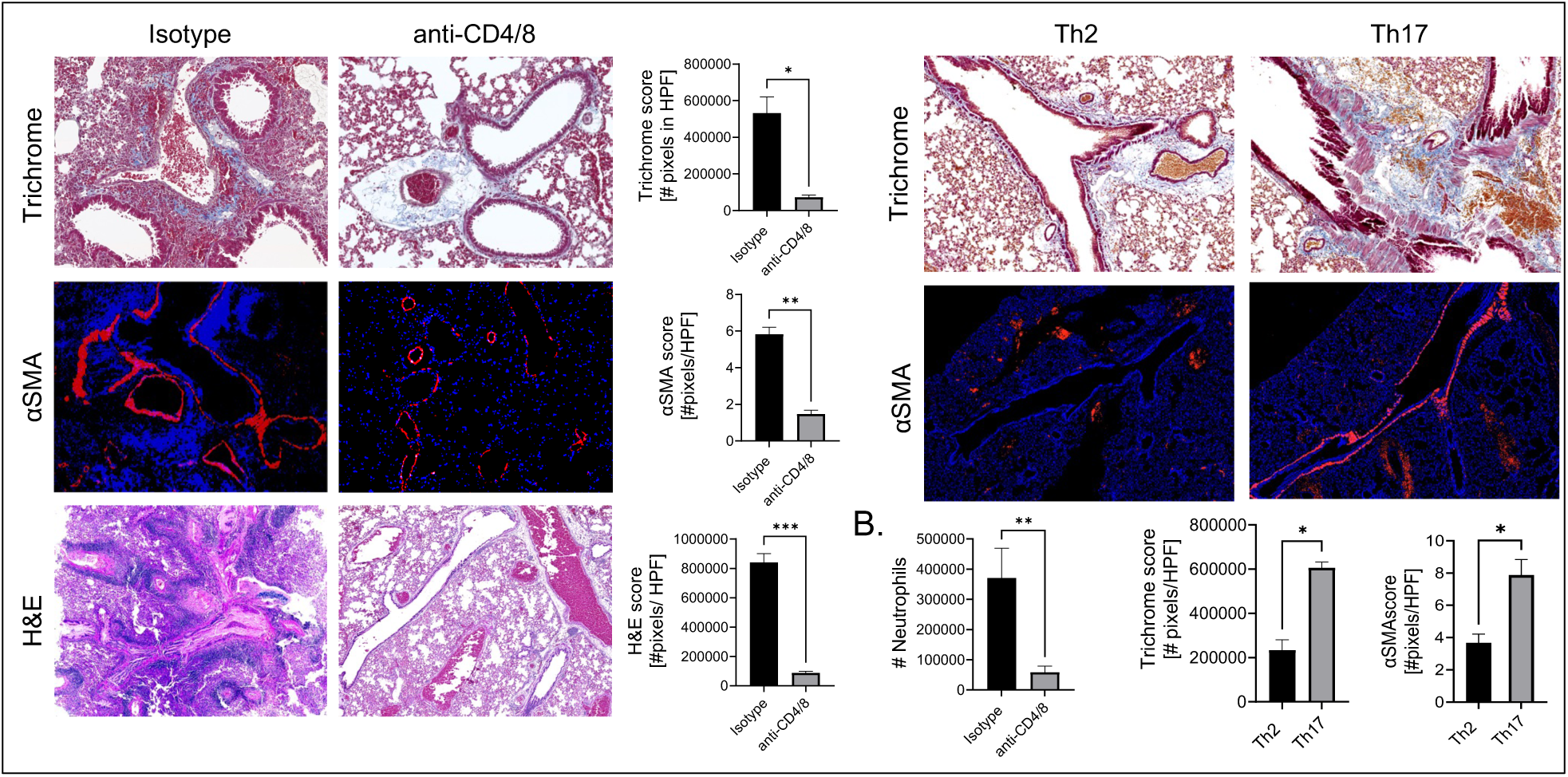
Th17 cells are essential initiators of MGA pathophysiology. (**A**) MGA-induced animals were treated with 100ug anti-CD4 and 100ug anti-CD8a or isotype control i.p. beginning on day 0 and continued every other day until harvest. Lung sections were stained and representative Masson trichrome (collagen), α-smooth muscle actin (αSMA), and hematoxylin and eosin (H&E) sections and accompanying quantifications are shown. (**B**) Number of neutrophils in the left lung lobe of T cell-depleted MGA induced mice, as assessed by flow cytometry. (**C**) 80,000 Th2 (CD45.2^+^CD3^+^CCR6^-^CCR4^+^CXCR3^-^CCR10^-^) or Th17 cells (CD45.2^+^CD3^+^CCR6^+^CCR4^+^CXCR3^-^CCR10^-^) were sorted from HDM challenged Balb/c or *Il4ra^-/-^* animals, respectively, and transferred intratracheally into naïve Rag1ko hosts. Recipient mice were challenged intranasally with 100ug HDM every other day for 2 weeks, followed by 2 consecutive challenges prior to harvest. Lung sections were stained and representative Masson trichrome and αSMA sections and accompanying quantifications are shown. Trichrome data show area of interest values covering full lung section of 4-8 individual mice per condition. αSMA data show mean values of all bronchi scored from 4-8 mice per condition. All data are representative of 2 independent experiments. *P < 0.05, **P < 0.005, ***P < 0.0005; HPF, high-power field.

### The mitochondrial OXPHOS pathway is enriched in MGA

Lung transcriptomic analyses revealed enrichment for expression of genes in the OXPHOS pathway in MGA mice (Fig 3A). We sorted 5 populations from MGA lungs: antigen-experienced T cells (CD45^+^CD3^+^CD44^+^CD62L^-^), eosinophils (CD45^+^Ly6C^+^SiglecF^+^), neutrophils (CD45^+^CD11b^+^GR1^+^SiglecF^-^), macrophages (CD45^+^CD11b^+^Ly6C^+^Mac3^+^auto-fluorescence^+^) and stromal cells (CD45^-^). Only T cells displayed an enrichment in mitochondrial OXPHOS gene expression (Fig. 3B), accounting for the MGA lung OXPHOS phenotype. *Il17* and *Rorc* transcripts were ∼112 and 62-fold-increased, respectively, in MGA T cells (Fig. 3B). We measured the rate of ATP production from mitochondrial OXPHOS in T cells sorted from MGA versus EA lungs. We observed a significant increase in oxygen consumption rate in MGA T cells (Fig. 3C), validating the transcriptomic analyses (Fig. 3B). Furthermore, we analyzed GEO dataset GSE89809 (43) comparing BAL CD3+ T cells of healthy versus severe asthmatics. We saw significant increase in an OXPHOS gene expression signature in severe asthmatic BAL T cells that coincided with increased IL-17-inducible chemokines and chemoattractants for neutrophils (43) (Fig. 3D). An additional study also showed non-Th2 asthmatic patients have high Th17 cells, and an increase transcript level of OXPHOS genes in their sputum (42).

**Figure 3:**
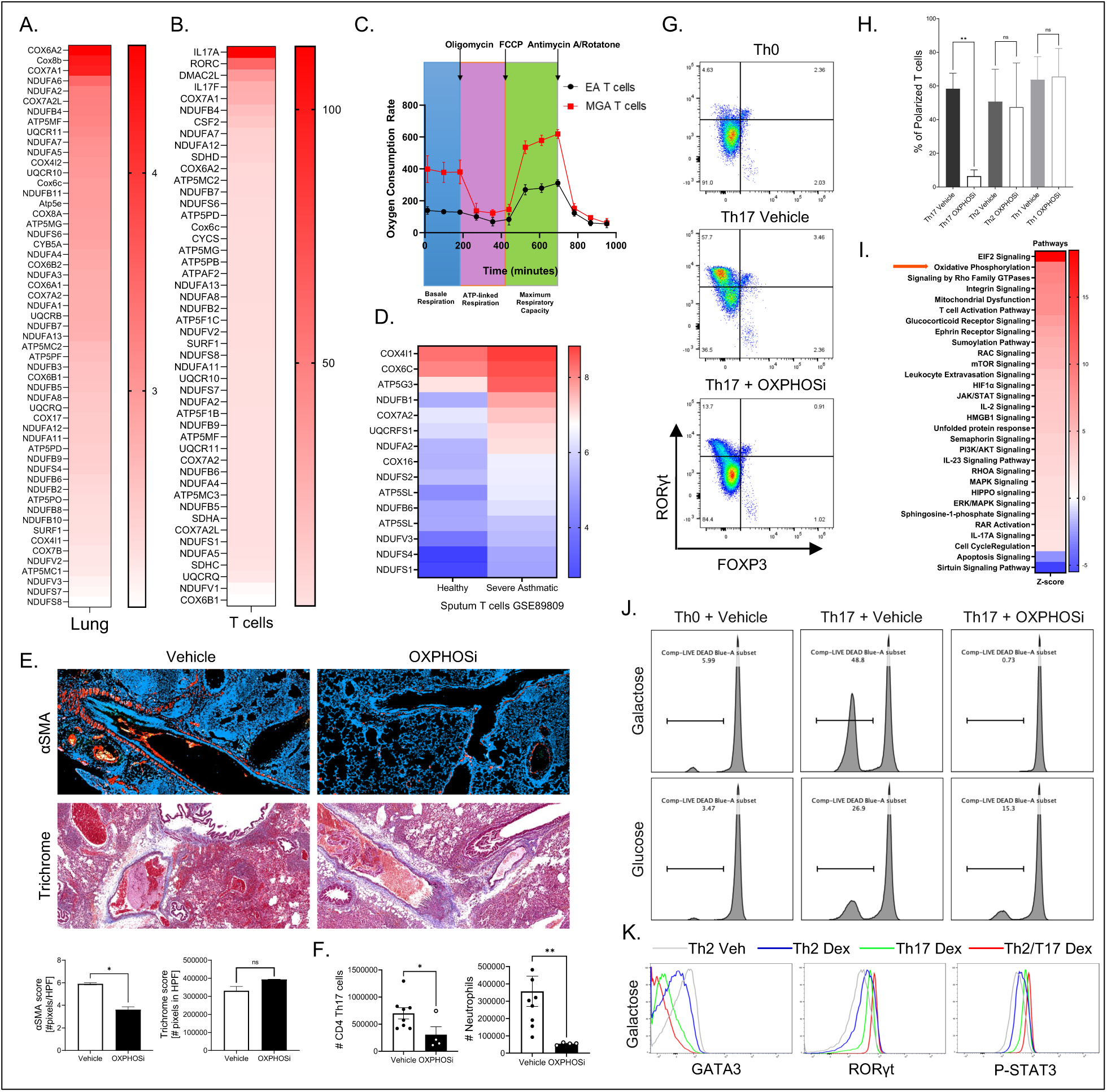
The mitochondrial OXPHOS pathway is enriched in MGA and OXPHOS controls Th17 cells via regulation of RORγt expression. (**A and B**) RNAseq from MGA versus EA bulk lung (A) or sorted CD45^+^CD3^+^CD44^+^CD62L^-^T effector cells (B) showing elevated OXPHOS signature. (**C**) Seahorse analysis to assess oxygen consumption rate for MGA versus EA T cells. **(D)** OXPHOS signature in human sputum T cells from severe asthmatics versus healthy controls, reanalyzed from GSE89809. (**E**) MGA-induced animals were treated with 150ug IACS-010759 i.p., versus vehicle control, beginning on day 7 and continuing every other day until harvest. Lung sections were stained and representative Masson trichrome and αSMA sections and accompanying quantifications are shown. Trichrome data show area of interest values covering full lung section of 4-8 individual mice per condition. αSMA data show mean values of all bronchi scored from 4-8 mice per condition. (**F**) Number of CD4 Th17 cells or neutrophils in the left lung lobe of MGA induced mice treated with or without OXPHOS inhibitor, as assessed by flow cytometry. (**G**) Naïve CD4 T cells were polarized under Th0 or Th17 conditions in the presence of vehicle or OXPHOS inhibitor for 72h. Representative plots are gated on live CD4 T cells and show expression of RORyt and Foxp3. Data are representative of 6 independent experiments. (**H**) Naïve CD4 T cells were polarized under Th1, Th2 or Th17 conditions in the presence of vehicle or OXPHOS inhibitor for 72h. Polarization percentages are shown. Data are representative of 6 independent experiments. (**I**) Pathway analysis of MGA versus EA T cells showing elevated OXPHOS signature. (**J**) Naïve CD4 T cells were polarized with Th17 conditions and vehicle or OXPHOS inhibitor in glucose-free RPMI supplemented with either 10mM galactose or 10mM glucose for 72h. Representative flow histogram of live cells expressed as percentage. Data are representative of 6 independent experiments. (**K**) Naïve CD4 T cells were polarized with Th2 conditions in glucose media or Th17 conditions in galactose media for 48h. Polarized Th2 cells, Th17 cells, or their co-culture (Th2/Th17) were treated with vehicle (Veh) or 5uM dexamethasone (Dex) in RPMI media (50:50 galactose and glucose mix) for 16h. Representative flow histograms of Gata3 (left), RORyt (middle), and pSTAT3 (right) of live CD4^+^CD44^+^CD62L^-^ T effector memory cells. Data are representative of 3 independent experiments. *P=0.05, **P < 0.005, HPF, high-power field; ns, nonsignificant.

### Inhibiting OXPHOS reduces inflammation

To inhibit OXPHOS, we treated *Il4rα^-/-^* at the challenge phase (day 7) post-disease establishment with a small drug inhibitor that binds complex I of the OXPHOS pathway (55) versus vehicle control. OXPHOS inhibition partially reduced airway remodeling. Smooth muscle hypertrophy was significantly reduced, whereas sub-epithelial collagen deposition was unchanged (Fig. 3E). Th17 cells and the associated neutrophilic inflammation were significantly decreased in MGA mice with OXPHOS inhibition (Fig. 3F).

### OXPHOS controls Th17 cells in MGA via regulation of RORγt expression

Using the OXPHOS inhibitor, we treated T cells polarized under Th1, Th2 and Th17 conditions in vitro. OXPHOS inhibition decreases RORγt expression only under Th17-polarizing conditions (Fig. 3G-H). Th1 and Th2 cells were not impacted by OXPHOS inhibition (Fig. 3H). GSEA analyses on MGA-sorted lung T cells show enrichment of EIF2 and mTOR signaling pathways, IL-17/-23 signaling, numerous pathways involved in TCR activation, and cell division, in addition to OXPHOS (Fig. 3I). Furthermore, transcripts of anti-apoptotic markers such as MCL1 are elevated and oxidative stress pathways, known for causing epithelial damage and airway remodeling, are also enhanced (HIF1α, HMGB1, UPR, etc.) (Fig. 3I). Interestingly, OXPHOS inhibition causes alteration in gene expression pathways for both positive and negative regulators of Th17 cells. These include the RhoA signaling pathway, which promotes Th17 cell differentiation (56), as well as the HIPPO pathway (57–59), and the Sirtuin signaling pathway, which impedes Th17 differentiation through STAT3 deacetylation (60) (Fig. 3I).

### OXPHOS controls Th17 cell survival

Naïve CD4 T cells were polarized with Th17 culture conditions and OXPHOS inhibitor or vehicle in glucose-free RPMI supplemented with either 10mM galactose, to promote OXPHOS, or 10mM glucose, to promote glycolysis. OXPHOS inhibition completely killed Th17 cells cultured in galactose medium, reversing their resistance to apoptosis as previously published (51), as opposed to glycolytic Th17 cells that exhibited ∼57% survival rate (Fig. 3J).

### Dexamethasone treatment skews Th2 cells towards Th17

Naïve CD4 T cells were polarized with Th2 conditions in glucose media or Th17 conditions in galactose media. Polarized Th2 cells, Th17 cells, or their co-cultures (Th2/Th17) were treated with vehicle (Veh) or 5uM dexamethasone (Dex) and stained for Th2 and Th17 markers. We show dexamethasone treatment downregulates GATA3 expression in Th2-exposed cells, reciprocal to an increase in RORγt and pSTAT3 expression (Fig. 3K). This is consistent with prior data showing glucocorticoids decrease IL-4Rα protein levels (53), in addition to dampening IL-5 and IL-13, and inducing apoptosis of Th2 cells.

### LIGHT controls airway remodeling in MGA

We analyzed the GEO dataset GSE137268 comparing EA versus MGA severe asthmatics and healthy controls. We observed a significant increase of *TNFSF14 (LIGHT)* transcripts in MGA asthmatics compared to EA or healthy controls (Fig. 4A). In HDM-induced *Il4ra^-/-^* mice, we observed a nearly 3-fold increase in lung transcript levels of *Tnfsf14* (Fig. 4B). We treated mice using LTβR-Fc, an antagonistic reagent that blocks LIGHT signaling with its receptors, starting at day 7. When LIGHT signaling was neutralized, the mice exhibit decreased sub-epithelial collagen deposition and smooth muscle hypertrophy (Fig. 4C). Reciprocally, intratracheal LIGHT injection promotes sub-epithelial collagen deposition and smooth muscle hypertrophy (Fig. 4D). We previously published LIGHT directly activates epithelial cells and fibroblasts, inducing their proliferation and production of pro-fibrotic molecules (44–47). Taken together, these data demonstrate that interrupting LIGHT signaling in severe asthma decreases pathological tissue remodeling.

**Figure 4:**
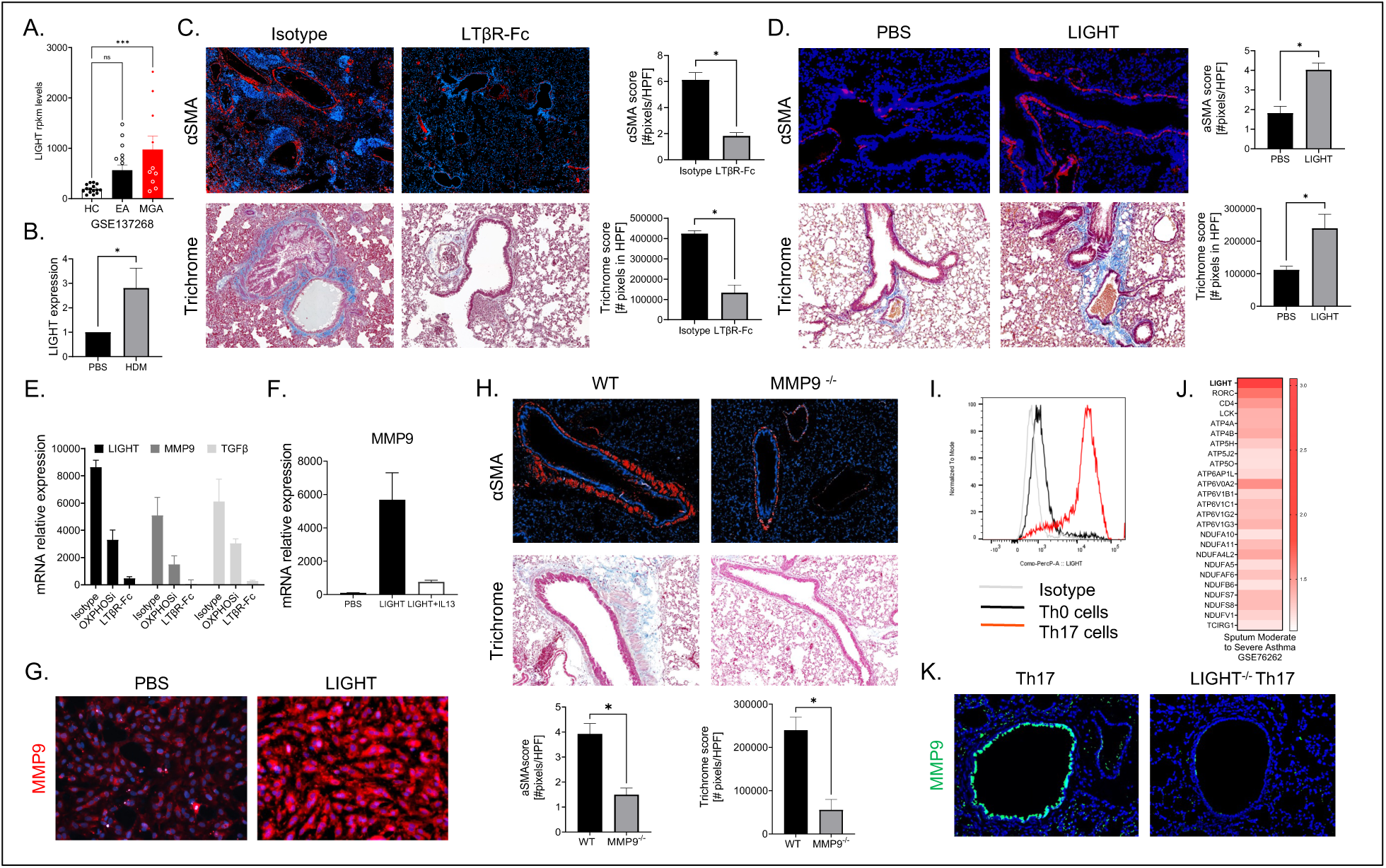
LIGHT controls airway remodeling in MGA via regulation of MMP9 expression. (**A**) LIGHT transcript levels (rpkm) in human MGA versus EA or healthy control counterparts, reanalyzed from GSE137268. (**B**) LIGHT expression in HDM-induced MGA mice versus PBS-induced control mice, represented as a ratio over PBS. (**C**) MGA-induced animals were treated with 100ug LTβR-Fc i.p., versus isotype control, beginning on day 7 and continuing every other day until harvest. Lung sections were stained and representative Masson trichrome and αSMA sections and accompanying quantifications are shown. (**D**) WT animals were given 10ug soluble rmLIGHT intratracheally for two consecutive days and harvested on day 3. Lung sections were stained and representative Masson trichrome and αSMA sections and accompanying quantifications are shown. (**E**) Quantitative PCR analysis was performed on lung samples from isotype, OXPHOS inhibitor or LTβR-Fc treated mice, and mRNA expression of LIGHT, MMP9, and TGF-β was calculated relative to L32. mRNA values are mean ± SEM of 4 mice per condition. (**F**). Normal Human Lung Epithelial Cells (NHBE cells) were stimulated with 10ng/mL rhLIGHT or rhLIGHT and 100ng/mL rhIL13 for 72h. Quantitative PCR analysis was performed on the NHBE and mRNA expression of MMP9 was calculated relative to L32. mRNA values are mean ± SEM of 4 mice per condition. (**G**) NHBE were stimulated with 10ng/mL rhLIGHT for 72h. Representative MMP9 staining is shown. (**H**) WT or *Mmp9^-/-^* animals were given 10ug soluble rmLIGHT intratracheally for two consecutive days and harvested on day 3. Lung sections were stained and representative Masson trichrome and αSMA sections and accompanying quantifications are shown. (**I**) Representative flow histogram of LIGHT for Th0 versus Th17 polarized cells. (**J**) LIGHT^high^ Th17 OXPHOS signature in sputum from human severe asthmatics versus healthy controls, reanalyzed from GSE76262. (**K**) Naïve CD4 T cells isolated from the lung of naïve *Il4ra*^- /-^ or *Il4ra^-/-^Tnfsf14^-/-^* mice were polarized with Th17 conditions in galactose media for 72h. 80k of these polarized Th17 or LIGHT^-/-^ Th17 cells were transferred intratracheally into naïve *Il4ra^-/-^Tnfsf14^-/-^* recipient hosts and harvested on day 4. Lung sections were stained and representative MMP9 staining is shown. Trichrome data show area of interest values covering full lung section of 4-8 individual mice per condition. αSMA data show mean values of all bronchi scored from 4-8 mice per condition. Data are representative of 2-3 independent experiments.*P < 0.05, ***P < 0.0005; HPF, high-power field.

### LIGHT controls airway remodeling in MGA via regulation of expression of MMP9

A remaining question is how LIGHT drives tissue remodeling post-allergen challenge in MGA. One possible mechanism is through MMP9. MMP9 can cleave TGFβ from its latent complex to activate the central fibrotic pathway and promote pathological remodeling (61). Additionally, MMP9 levels are dramatically increased in acute asthmatics (62) and severe patients with high asthma exacerbation rates (63). Our data show that human bronchial epithelial cells stimulated with LIGHT enhance MMP9 transcripts by qPCR analysis (Fig. 4F) and MMP9 protein levels (Fig. 4G). Furthermore, LIGHT pro-fibrotic activity in the lung is fully dependent on TGFβ, and not IL-13, since LIGHT-induced wildtype mice treated with anti-TGFβ are completely protected from airway remodeling (46). Our data demonstrate a similar trend of LIGHT, MMP9, and TGFβ expressions in the lungs of mice with MGA, correlating with fibrosis severity (Fig. 4E-F). Furthermore, LIGHT-induced *Mmp9^-/-^* mice dramatically decreased α-smooth muscle hypertrophy, in the absence of exogenous TGFβ, compared to wildtype mice (Fig. 4H). Lastly, LIGHT-induced MMP9 expression on epithelial cells decreased in the presence of IL-13 (Fig. 4F). This finding could be a decisive factor between the mild steroid-sensitive asthma endotype (eosinophilic Th2) and the severe steroid resistant one (neutrophilic Th17) and would explain the enhanced airway remodeling seen in MGA compared to EA.

### T cells are the main cellular sources producing LIGHT in MGA

Flow cytometry analysis confirmed an increase in surface expression of LIGHT in Th17 cells (Fig. 4I) compared to Th0 cells. Furthermore, we analyzed the GEO dataset GSE76262 and showed LIGHT^high^ T cells express enhanced Th17 and OXPHOS gene expression signature in the sputum of human severe asthmatics compared to healthy controls (Fig. 4J). Naïve CD4 T cells isolated from the lungs of *Il4ra^-/-^* or *Il4ra^-/-^Tnfsf14^-/-^* mice were polarized with Th17 conditions in galactose media for 72h. Equivalent numbers of Th17 versus LIGHT^-/-^ Th17 cells were transferred intratracheally into naïve *Il4ra^-/-^Tnfsf14^-/-^* recipient hosts and harvested on day 4. In lung biopsies from mice that received Th17 cells lacking LIGHT, there was a striking decrease in MMP9 expression by bronchial epithelial cells (Fig. 4K). This suggests Th17 cells are the main cellular source of LIGHT that drives fibrosis.

### LIGHT and OXPHOS synergize to control MGA

Aside from producing LIGHT, OXPHOS high Th17 cells induce dynamic processes of airway remodeling involving fibroblast recruitment and activation. One insight into how OXPHOS promotes airway remodeling came from our data showing MGA-sorted Th17 cells significantly upregulate Osteopontin (OPN; >307 fold) (Fig.5A), which is elevated in human severe asthmatics (64–66). OPN is an extracellular matrix glycoprotein that also serves as a cytokine, bridging inflammation with tissue repair in lung diseases through control of fibroblast activation (67, 68). Lung fibroblasts cultured with Th0 or Th17 polarized cells, increase their expression of α-smooth muscle actin, which is abrogated when anti-OPN neutralizing antibody is added (Fig.5B). This suggests that Th17-derived OPN activates fibroblasts to promote smooth muscle hypertrophy. Furthermore, the transfer of LIGHT^-/-^ Th17 cells into *Il4ra^-/-^Tnfsf14^-/-^* mice resulted in decreased OPN levels compared to mice receiving wildtype Th17 cells, although not fully abrogated (Fig. 5C). Thus, OPN expression, which is a product of Th17 cells in MGA responsible for fibroblasts activation and myofibroblast differentiation, is partially dependent on LIGHT.

**Figure 5:**
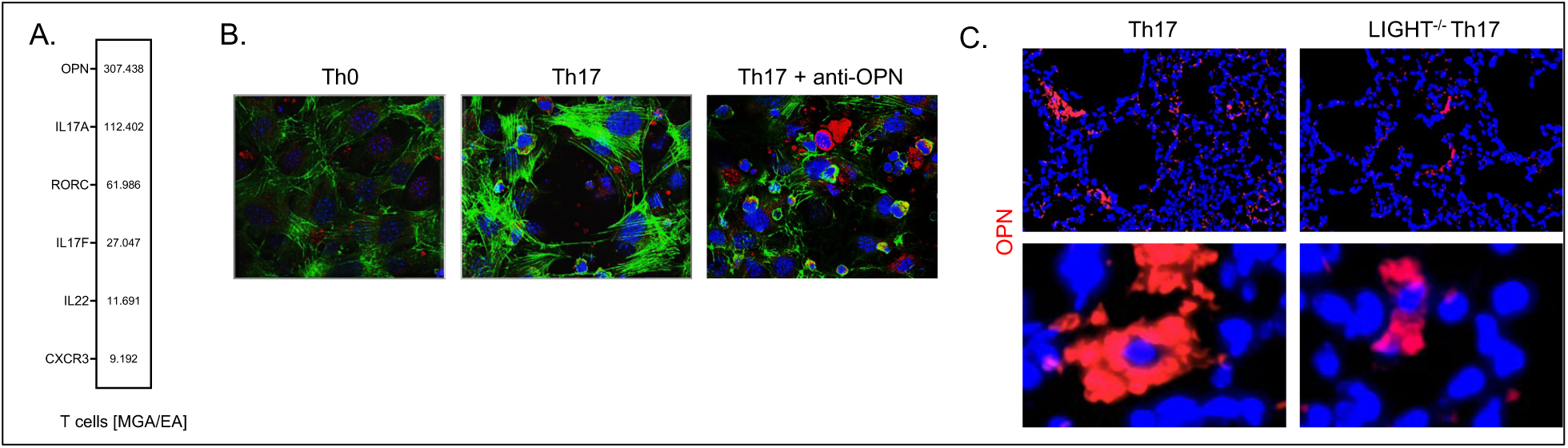
OXPHOS regulates Th17-derived OPN to drive fibroblast activation. (**A**) RNAseq from MGA versus EA sorted CD45^+^CD3^+^CD4^+^CD44^+^CD62L^-^ T effector cells showing elevated OPN-Th17 signature. (**B**) Primary WT mouse fibroblasts (50,000) were co cultured with 1 million naïve CD4 T cells in DMEM media containing Th0 control or Th17 polarizing conditions, with or without 2ug/mL anti-OPN neutralizing antibody for 72h. Co-cultures were stained for Phalloidin and DAPI nuclear stain with representative 60X images are shown. (**C**) Naïve CD4 T cells isolated from the lung of naïve *Il4ra^-/-^* or *Il4ra^-/-^Tnfsf14^-/-^* mice were polarized with Th17 conditions in galactose media for 72h. 80k of these polarized Th17 or LIGHT^-/-^ Th17 cells were transferred intratracheally into naïve *Il4ra^-/-^Tnfsf14^-/-^* recipient hosts and harvested on day 4. Lung sections were stained and representative OPN staining is shown. Data are all representative of 2-3 independent experiments.

### LIGHT and OXPHOS dual blockade abrogates severe asthma and increases the frequency of suppressive Tregs

We compared antagonistic blockade of LIGHT or OXPHOS alone to combination therapy neutralizing both LIGHT and OXPHOS simultaneously. All treatments were given starting day 7, after disease establishment. Inhibiting LIGHT significantly reversed sub-epithelial collagen deposition and airway fibrosis (Fig. 6A) with a mild decrease in airway neutrophilia (Fig. 6C). Vice versa, OXPHOS inhibition dramatically decreased Th17 cells and airway neutrophilia, without improving collagen deposition. Importantly, LIGHT and OXPHOS co-blockade completely resolved airway remodeling, Th17 cellularity, and airway neutrophilia (Fig. 6 A-C). The combination therapy dampened airway hyperreactivity in response to methacholine (Fig. 6D) to a greater extent than either monotherapy (Fig. 6D). Furthermore, reciprocal to the decrease in Th17/neutrophilic inflammation, we observed an increase in Foxp3^+^CD25^+^CD127^Low^ Treg cells in the combined anti-OXPHOS/LIGHT treatment (Fig. 6B). This was not seen in the monotherapy LTβR-Fc or OXPHOS inhibitor treated groups (Fig. 6B).

**Figure 6:**
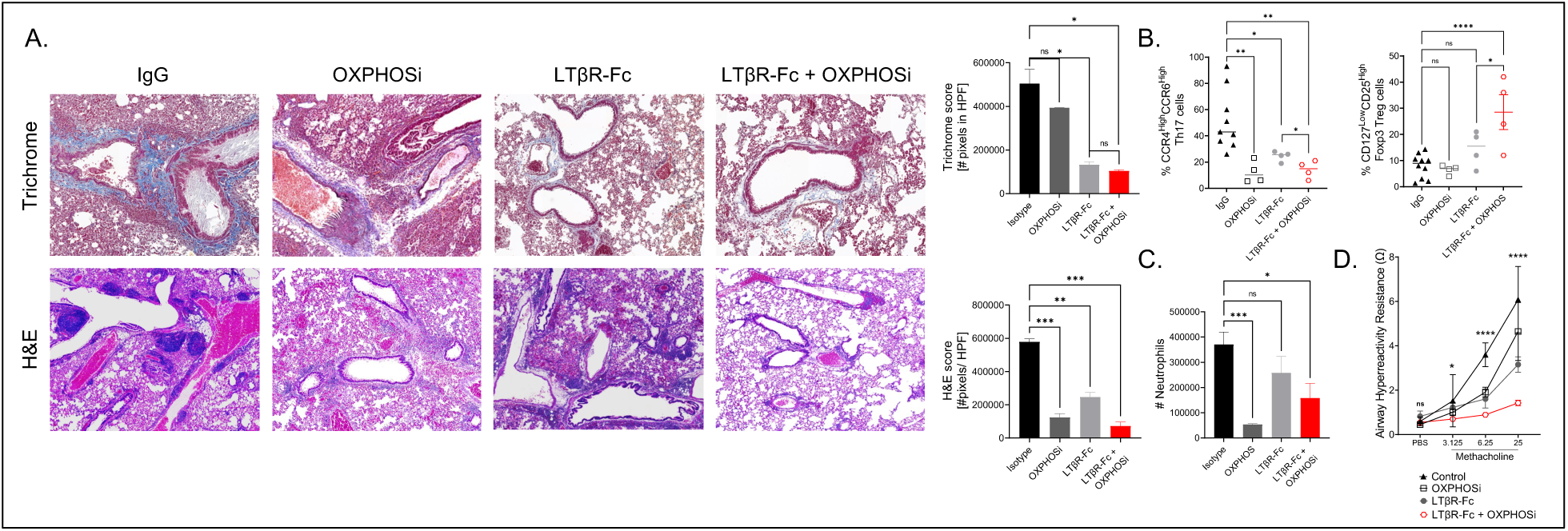
LIGHT and OXPHOS dual blockade abrogates severe asthma. (**A**) MGA-induced animals were treated with isotype control, 100ug LTβR-Fc, 150ug Oxphos inhibitor (IACS-010759), or LTβR-Fc + Oxphos inhibitor beginning on day 7 and continuing every other day until harvest. Lung sections were stained and representative Masson trichrome and αSMA sections and accompanying quantifications are shown. (**B**) Percentage of CCR4^high^CCR6^high^ Th17 cells and CD127^low^CD25^high^ Foxp3 Treg cells by flow cytometry from left lung lobe of isotype, Oxphos inhibitor, LTβR-Fc or combination (LTβR-Fc + Oxphosi) treated MGA induced mice. (**C**) Number of neutrophils in the left lung lobe of MGA induced mice treated with isotype, Oxphos inhibitor, LTβR-Fc or combination (LTβR-Fc + Oxphosi), as assessed by flow cytometry. (**D**) Airway resistance to increasing doses of methacholine exposure (4-6 mice per group). Trichrome data show area of interest values covering full lung section of 6 individual mice per condition. αSMA data show mean values of all bronchi scored from 6 mice per condition. Data are representative of 3 independent experiments. *P < 0.05, **P < 0.005, ***P < 0.0005, ****P < 0.00005; HPF, high-power field; ns, nonsignificant.

### LIGHT/OXPHOS signature is enhanced in multiple Th17-fibrotic diseases

GEO datasets containing human Th17 fibrotic disease cohorts were analyzed to show elevated LIGHT/OXPHOS gene signatures, including idiopathic pulmonary fibrosis (IPF) (Fig. 7A and B chronic obstructive pulmonary disease (COPD) (Fig. 7C), Crohn’s disease (CD) (Fig. 7D), rheumatoid arthritis (RA) (Fig. 7E), psoriasis (Pso) (Fig. 7F), psoriatic arthritis (PsA) (Fig. 7G), systemic lupus erythematosus (SLE) (Fig. 7H), and multiple sclerosis (MS) (Fig. 7I).

**Figure 7:**
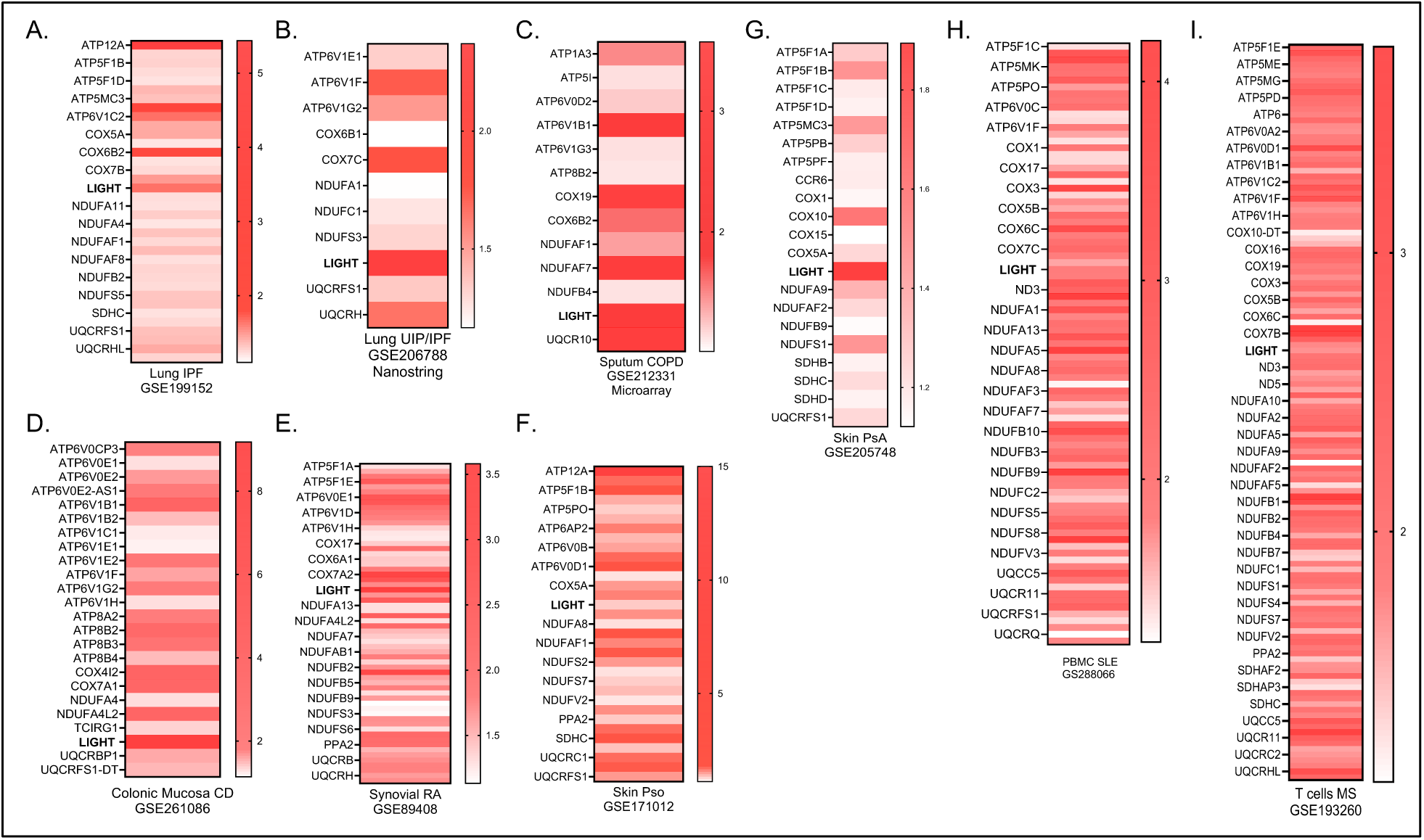
LIGHT and OXPHOS transcripts are elevated in human patients with Th17 fibrotic diseases. GSE datasets containing human Th17 fibrotic disease cohorts were reanalyzed to show elevated LIGHT-OXPHOS signatures, including (**A and B**) idiopathic pulmonary fibrosis (IPF), (**C**) chronic obstructive pulmonary disease (COPD), (**D**) Crohn’s disease (CD), (**E**) rheumatoid arthritis (RA), (**F**) psoriasis (Pso), (**G**) psoriatic arthritis (PsA), (**H**) systemic lupus erythematosus (SLE), and (**I**) multiple sclerosis (MS).

## DISCUSSION

Despite its enrichment in the lungs and bronchial lavage of severe asthmatics, targeting IL-17 signaling using brodalumab (an IL-17 receptor antagonist) showed no clinical response nor statistically significant improvements in primary and secondary endpoints (54). This surprising failure can be attributed to: 1) a poor selection of the patient cohort, with no prior evidence of high sputum IL-17 levels; 2) a poor timing of IL-17 targeting; 3) a highly unlikely off-target effect; 4) the need to co-block Th2 cytokines in addition to IL-17 (this is also unlikely since severe asthmatics have airways enriched with Th17 cells, and have already received high doses of corticosteroids that could kill Th2-driven eosinophilia). Moreover, in a phase I trial, the humanized IgG4 bispecific antibody targeting both IL-13 and IL-17 was well-tolerated in healthy volunteers, but anti-drug antibodies were observed (69); 5) a poor ability of IL-17 to directly drive pathological airway remodeling in structural cells (lung epithelial cells, smooth muscle cells, and fibroblasts). Indeed, compared to the central fibrotic cytokine TGFβ, IL-17 direct stimulation of lung fibroblasts drove weak smooth muscle hypertrophy and collagen deposition, and minimal epithelial to mesenchymal transition (70, 71). Thus, fundamental gaps in knowledge include how Th17 cells induce pathological remodeling independently of IL-17 and how Th17 differentiation occurs in MGA.

Existing MGA models have been reported, using cyclic-di-GMP combined with HDM (cyclic-di-GMP/HDM) challenges performed on wildtype (IL4Rα-sufficient) mice (72). However, this generated a mixed Th1/Th2^Low^/Th17 response, which differs from human severe asthmatics with OXPHOS^high^Th17^high^ phenotype (42). Notably, the cyclic-di-GMP/HDM model is Th1 dominant (72); it stalled at demonstrating an upregulation of OXPHOS seen in human non-Th2 severe asthma, likely due to a mixed Th1/Th17 phenotype. No airway remodeling was shown either in this cyclic-di-GMP/HDM model (72). While we do not fully understand why some asthmatic patients fail to mount a proper Th2 response resulting in MGA development, we know steroids kill Th2 cells and eosinophils, but neutrophils and Th17 cells are resistant (26). Moreover, glucocorticoid downregulates levels of IL4Rα (53), in addition to dampening IL-5 and IL-13, and inducing apoptosis of Th2 cells (73). Compared to Th2 cells, Th17 cells are not sensitive to dexamethasone in vitro or in vivo, mediating steroid-resistance (26). Thus, prolonged treatment to steroids in patients can eradicate the Th2 response in their lungs while sustaining a resistant Th17 response, making them Th2-low. These Th2-low patients have little to no functional IL4Rα signaling, reciprocal to a dominant Th17 response resistant to apoptosis.

To accurately model these Th2-low patients in mice, and overcome the significant limitation imposed by Th2 responses in the previously published model (72), we genetically ablated *Ilr4α* to impose a defect in IL4/IL13 signaling. This model drives significant airway remodeling with concomitant increase in pathogenic Th17 cells that are OXPHOS^high^ mimicking the non-Th2 asthmatic patients. We confirmed that this model is steroid-resistant and does not respond to anti-IL17 immunotherapy, similarly to human MGA. We show T cells are essential drivers of MGA. MGA-sorted Th17 cells promote greater airway remodeling than EA-sorted Th2 cells, supporting their enhanced pathogenicity. Furthermore, we demonstrate that mitochondrial OXPHOS is essential for generation of pathogenic Th17 cells. OXPHOS^high^ T cells enrich pathways involved in Th17 development like IL-23 signaling, RHOA signaling, HIPPO signaling, and IL-17 itself while dampening negative regulators like the sirtuin signaling pathway. This aligns with previous work showing OXPHOS plays a pivotal role for lineage specification towards pathogenic Th17 differentiation by regulating BATF (52). We show OXPHOS regulates RORγt expression and Th17 programming, dysregulating the Th17/Treg balance in favor of Type 3 inflammation. OXPHOS inhibition during differentiation is sufficient to abrogate RORγt^+^Th17 cells, both in vitro and in vivo. Previous work showed OXPHOS promotes anti-apoptotic survival of Th17 cells marked by high BCL-XL and low BIM. We observed a survival advantage of OXPHOS-positive Th17 cells, likely due to enhanced anti-apoptotic markers like MCL1. Conversely, we show that OXPHOS inhibition kills Th17 cells cultured under OXPHOS conditions (galactose) but not in glycolytic media (glucose). Th1 and Th2 cells remain insensitive to OXPHOS inhibition. Moreover, Th2 cells exposed to dexamethasone downregulate GATA3 and Th2 markers, reciprocal to increases in RORγt and pSTAT3. This explains why MGA patients accumulate Th17 cells in their lungs, driven by skewing the inflammatory endotype towards pathogenic Th17 cells that increase OXPHOS to acquire resistance to apoptosis. The transfer of allergic OXPHOS-positive Th17 cells exacerbates airway remodeling compared to allergic Th2 cells. This elucidates why MGA patients have greater lung fibrosis than EA patients and emphasizes the need for anti-fibrotics, rather than anti-inflammatory therapeutics like inhaled corticosteroids. This positions severe asthma among the “fibrotic” diseases rather than solely considering it an inflammatory disorder. We have evidence that OXPHOS inhibition not only decreased Th17 cellular development during polarization but also effectively killed them, restoring apoptotic capacity and sensitivity to steroids. This is important as OXPHOS inhibition could reverse corticosteroid resistance seen in MGA patients. Targeting mitochondrial OXPHOS has been extensively investigated in cancer, and other hypoxic diseases. For MGA patients, OXPHOS inhibition offers a potential therapeutic avenue for those with early diagnosis, when profibrotic LIGHT is not yet made. If the disease is advanced, killing Th17 cells may not suffice to reverse fibrosis driven by downstream molecules like LIGHT and OPN.

In vivo, OXPHOS inhibition post-disease onset led to significant decrease in Th17 cells and subsequent airway neutrophilia, in addition to α-smooth muscle actin hypertrophy. However, pre-formed collagen was maintained due to early expression of LIGHT by Th17 cells during disease initiation. Depletion of T cells during disease initiation fully abolished LIGHT expression and fibrosis, with dramatic decrease in collagen and α-smooth muscle actin accumulation. Neutralizing LIGHT post-disease onset decreased fibrosis, albeit to a lesser extent than T cell depletion, but does not kill Th17 cells on its own, so disease might reoccur in chronically exposed allergic patients. LIGHT promotes airway remodeling through upregulation of MMP9, a molecule involved in releasing active TGFβ from its latent form leading to fibrosis. Recombinant LIGHT injection in the lungs fails to promote airway remodeling in *Mmp9^-/-^* compared to wildtype controls. LIGHT directly stimulates MMP9 expression by bronchial epithelial cells, and Th17 cells are the primary source of LIGHT driving this activity. Moreover, Th17 cells amplify the fibrotic response via expression of OPN, a molecule involved in fibroblasts’ activation, in a partially LIGHT-dependent manner. Human severe asthma patients with high exacerbation rates upregulate MMP9 (63) and OPN (64–66). Excitingly, LIGHT and OXPHOS concomitant neutralization, post-disease onset, completely eradicates all features of disease, killing pathogenic Th17 cells and reversing preformed collagen and smooth muscle actin. These findings offer a novel therapeutic approach to treat severe chronic disorders presenting with fibrosis and pathogenic Th17 cell involvement. While neutralizing IL-17 itself did not reverse disease, eliminating Th17 cells through OXPHOS inhibition combined with LIGHT blockade, abolished all features of disease.

Here, we have a first line of evidence showing that targeting LIGHT and OXPHOS is effective in treating fibrotic diseases involving pathogenic Th17 cells. This includes severe asthma but also extends to multiple sclerosis, rheumatoid arthritis, chronic obstructive pulmonary disease, psoriasis, psoriatic arthritis, systemic lupus, and idiopathic pulmonary fibrosis, in which we found a dominant LIGHT/OXPHOS gene signature. Thus, understanding the metabolic reprogramming of Th17 cells promoting severe fibrosis and this combination therapy offers tremendous therapeutic benefits to human health.

## MATERIALS AND METHODS

### Mice

Six- to eight-week-old female and male WT Balb/c (JAX:000651), *Ilr4a^-/-^* mice (JAX:003514), *Mmp9^-/-^* mice (JAX:007084), Rag1^-/-^ mice (JAX:003145), *Tnfsf14^-/-^*(generous gift from Dr Klasus Pfeffer, Dusseldorf University, Germany)(74), and *Ilr4a^-/-^Tnfsf14^-/-^* (generated in-house) mice were purchased directly from JAX or bred in house. In all studies, 4- to 8-week-old male and female mice were used. All studies and protocols were approved by and in compliance with the regulations of the Institutional Animal Care and Use Committees of Cincinnati Children’s Hospital Medical Center.

### Experimental in vivo protocols

To model EA or MGA, Balb/c or *Ilr4a^-/-^* mice were used respectively. Mice were given 100ug house dust mite (HDM; Greer) in 20uL PBS intranasally for three consecutive weeks (five consecutive days of induction followed by two days resting) with a final dosage of intranasal HDM given 24 hours prior to harvest. For dexamethasone treatment, HDM induced *Ilr4a^-/-^* mice were treated with 50ug dexamethasone (Sigma) intratracheally/mouse one hour prior to allergen challenge beginning day 7 and continued through the end of the experiment. To block IL-17 signaling, 100ug anti-IL17A (17F3; BioXCell) or matching isotype control (MOPC-21; BioXCell) were given intraperitoneally (i.p.) to HDM induced *Ilr4a^-/-^* mice beginning on day 7 and continuing every other day for a total of 7 injections. To inhibit OXPHOS, the small drug inhibitor IACS-010759 (Cayman Chemical) was used. HDM induced *Ilr4a^-/-^* mice were treated with 150ug IACS-010759 i.p., versus vehicle control, beginning on day 7 and continuing every other day for a total of 7 injections. For neutralizing LIGHT, HDM induced *Ilr4a^-/-^* mice were treated with 100ug i.p. LTβR-Fc (Elabscience), an antagonistic reagent that blocks LIGHT signaling with its receptors, or matching isotype control starting at day 7 and continuing every other day until for a total of 7 injections. For LIGHT-induction, mice were given 10ug recombinant mouse LIGHT (R&D) or PBS intratracheally for two consecutive days and sacrificed on day 3. For T cell depletion, 100ug anti-CD4 (GK1.5; BioXCell) and 100ug anti-CD8a (YTS 169.4; BioXCell) or matching isotype control (LTF-2 Rat IgG2b; BioXCell) were given i.p. beginning on day 0 prior to HDM induction and continuing every other day until the end of the experiment for a total of 10 doses.

For Th2 or Th17 transfer studies in vivo, Balb/c or *Il4ra^-/^*^-^ animals were given 5 consecutive doses of 100ug HDM intranasally. Lungs were processed and T cells were sorted using negative selection magnetic sorting. Briefly, cells were incubated with 1:500 biotinylated antibodies (CD11b, CD11c, CD45R, CD49b, IgG1k; Invitrogen) for 30 minutes on ice, followed by a PBS rinse and subsequent incubation with 1:10 diluted anti-biotin microbeads (Miltenyi Biotec) for 30 minutes in FACS tubes. Using magnetic separator, negative fraction was removed and passed through three sequential rounds of separation. Sorted T cells were incubated with flow antibodies to CD45.2, CD3, CCR4, CXCR3, CCR10, and CCR6, before being resuspended in 7-AAD (Biolegend) and sort buffer (1X PBS Ca^2+^/Mg^2+^-free, 2% FBS, and 25mM Hepes). Cells were sorted using FACSAriaFusion (BD) for the following populations: live CD45.2^+^CD3^+^CCR6^-^CCR4^+^CXCR3^-^CCR10^-^ Th2 cells from Balb/c pre-sorted T cells or live CD45.2^+^CD3^+^CCR6^+^CCR4^+^CXCR3^-^CCR10^-^ Th17 cells from *Il4ra^-/^*^-^ pre-sorted T cells. For transfer in vivo, 80k Th2 or Th17 cells were mixed with 100ug HDM and transferred intratracheally into naïve Rag1ko recipient hosts. On day 3, the recipient mice were challenged intranasally with 100ug HDM every other day for 2 weeks, followed by 2 consecutive challenges prior to harvest (for a total of 8 intranasal injections).

For transfer of Th17 or LIGHT^-/-^ Th17 cells in vivo, T cells were isolated from the lung of naïve *Il4ra^-/^*^-^ or *Ilr4a^-/-^Tnfsf14^-/-^* by negative selection magnetic sorting (as described above). Isolated T cells were cultured in glucose-free RMPI (Gibco) supplemented with 10% FBS, 1% PSG and 10mM galactose (Sigma) and were incubated with 5ug/mL bound anti-CD3 (clone 145-2C11; Biolegend), 2ug/mL anti-CD28 (clone 37.51; Tonbo) for 2h at 37C before adding polarizing conditions. For Th17 polarization, T cells were cultured with 40ng/ml IL-6 (PeproTech), 40ng/ml IL-23 (Biolegend), 0.5ng/ml TGFB (R&D), 10ug/mL anti-IL-4 (clone 11B11; BioXCell), and 10ug/mL anti-IFNy (clone H22; BioXCell) in RPMI for 72h at 37C. Polarized cells were collected, and dead cells were removed via Percoll gradient. For transfer in vivo, 80k Th17 or LIGHT^-/-^ Th17 cells were transferred intratracheally into naïve *Ilr4a^-/-^Tnfsf14^-/-^* recipient hosts and harvested on day 4.

### Assessment of AHR

Mice were anesthetized with ketamine/xylazine (each at 100mg/kg) delivered i.p. and tracheostomized, which was followed by delivery of pancuronium bromide i.p. (30ug/injection). Airway resistance was then measured using the FlexiVent system (Flexiware v8.2; SCIREQ). Mice were mechanically ventilated at 150 breaths/min with positive end-expiratory pressure (PEEP) of 3 cm H2O and were given time to acclimate. Airway hyperresponsiveness (AHR) was assessed by challenge of increasing doses of aerosolized methacholine (MCh; 0, 3.125, 6.25, 25mg/ml; Sigma) which were delivered for 10 seconds each, followed by measurement of airway resistance.

### Assessment of pulmonary inflammation by flow cytometry

Lungs were dissociated as previously described(75). Cells were stained with monoclonal antibodies to for flow cytometry to assess immune cell populations (see Table 1). For intracellular staining, the Cytofix/Cytoperm kit (BD Biosciences) was used for fixation and permeabilization. Flow analysis was done on either a Fortessa device (BD Biosciences) or an Aurora (Cytek). All data were analyzed using FlowJo v.10.8.1 software.

**Table 1:**
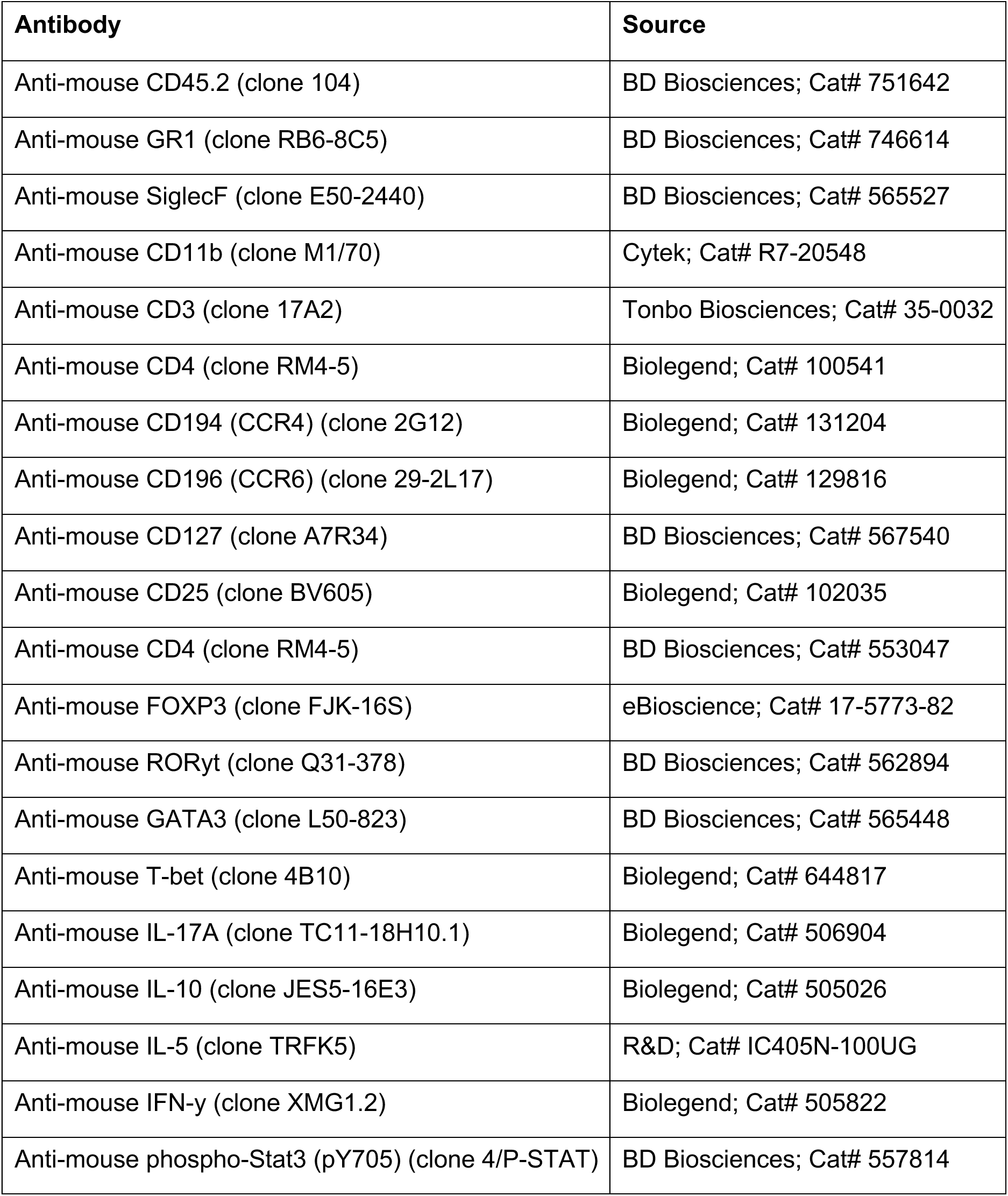
Flow cytometry antibodies. A list of flow cytometry antibodies and their clones used.

| Antibody | Source |
| --- | --- |
| Anti-mouse CD45.2 (clone 104) | BD Biosciences; Cat# 751642 |
| Anti-mouse GR1 (clone RB6-8C5) | BD Biosciences; Cat# 746614 |
| Anti-mouse SiglecF (clone E50-2440) | BD Biosciences; Cat# 565527 |
| Anti-mouse CD11b (clone M1/70) | Cytek; Cat# R7-20548 |
| Anti-mouse CD3 (clone 17A2) | Tonbo Biosciences; Cat# 35-0032 |
| Anti-mouse CD4 (clone RM4-5) | Biolegend; Cat# 100541 |
| Anti-mouse CD194 (CCR4) (clone 2G12) | Biolegend; Cat# 131204 |
| Anti-mouse CD196 (CCR6) (clone 29-2L17) | Biolegend; Cat# 129816 |
| Anti-mouse CD127 (clone A7R34) | BD Biosciences; Cat# 567540 |
| Anti-mouse CD25 (clone BV605) | Biolegend; Cat# 102035 |
| Anti-mouse CD4 (clone RM4-5) | BD Biosciences; Cat# 553047 |
| Anti-mouse FOXP3 (clone FJK-16S) | eBioscience; Cat# 17-5773-82 |
| Anti-mouse RORyt (clone Q31-378) | BD Biosciences; Cat# 562894 |
| Anti-mouse GATA3 (clone L50-823) | BD Biosciences; Cat# 565448 |
| Anti-mouse T-bet (clone 4B10) | Biolegend; Cat# 644817 |
| Anti-mouse IL-17A (clone TC11-18H10.1) | Biolegend; Cat# 506904 |
| Anti-mouse IL-10 (clone JES5-16E3) | Biolegend; Cat# 505026 |
| Anti-mouse IL-5 (clone TRFK5) | R&D; Cat# IC405N-100UG |
| Anti-mouse IFN-γ (clone XMG1.2) | Biolegend; Cat# 505822 |
| Anti-mouse phospho-Stat3 (pY705) (clone 4/P-STAT) | BD Biosciences; Cat# 557814 |

### Immunohistochemistry and immunofluorescence microscopy on lung sections

Hematoxylin and Eosin (H&E) and Masson Trichrome staining followed manufacturer recommendations (Poly Scientific R&D and Thermo Fisher Scientific) and slides were mounted using Cytoseal 60. Sections were scanned with the Zeiss Axioscanner (20X) and analyzed using ImagePro Premier. To assess smooth muscle actin (aSMA), OPN, or MMP9 staining on paraffin-embedded tissues, deparaffinization was performed by incubation in xylene and subsequent incubations in ethanol, followed by PBS rinse. Slides were then heated for 20 minutes in antigen retrieval solution (citrate buffer pH 6). After a PBS wash, sections were incubated in a blocking solution with 10% by volume skim milk powder, 10% donkey serum and 1% Fc block for 1 hour at RT. Primary antibody for aSMA (1:350, ab5694; Abcam), OPN (1:400, MAB808-SP; R&D), or MMP9 (1:400, ab76003; Abcam) was added overnight at room temperature. For the fluorescent signal, donkey anti-rabbit Rhodamine RedX secondary antibody (ImmunoJackson Research) or donkey anti-rabbit AF488 secondary antibody (ImmunoJackson Research) is added at 1:500 dilution for 3 hours at room temperature. Nuclear stain is performed by using DAPI at 1:5000 dilution for 5 minutes at room temperature. Sections were mounted with Cytoseal 60, scanned 24h later with the Zeiss Axioscanner (20X) and analyzed using ImagePro Premier.

### In vitro T cell polarization and analysis

T cells were isolated from lung and spleen of 4-8wk old naive WT mice using negative selection magnetic sorting. Briefly, cells were incubated with biotinylated antibodies (CD11b, CD11c, CD45R, CD49b, IgG1k; Invitrogen) 1:500 for 30 minutes on ice, followed by PBS rinse and subsequent incubation with 1:10 diluted anti-biotin microbeads (Miltenyi Biotec) for 30 minutes in FACS tubes. Using magnetic separator, negative fraction was removed and passed through three sequential rounds of separation. Isolated T cells were cultured in RPMI 1640 (Gibco) supplemented with 10% heat inactivated fetal bovine serum (FBS) (Corning) and 1% Penicillin-Streptomycin-Glutamine (PSG) (Gibco) or glucose-free RMPI (Gibco) supplemented with 10% FBS, 1% PSG and 10mM galactose (Sigma). T cells were incubated with 5ug/mL bound anti-CD3 (Biolegend) and 2ug/mL anti-CD28 (Tonbo) for 2h at 37C before adding polarizing conditions. For Th17 conditions, T cells were cultured with 40ng/ml IL-6 (PeproTech), 40ng/ml IL-23 (Biolegend), 0.5ng/ml TGFB (R&D), 10ug/mL anti-IL-4 (BioXCell), and 10ug/mL anti-IFNy (BioXCell) in RPMI for 72h at 37C. For Th2 conditions, T cells were cultured with 50ng/mL IL-4 (PeproTech), 20ng/mL IL-2 (R&D), and 10ug/mL anti-IFNy in RPMI for 72h at 37C. For Th1 conditions, T cells were cultured with 50ug/mL IFNy (PeproTech), 10ng/mL IL-12 (Peprotech), 20ng/mL IL-2 (R&D), and 10ug/mL anti-IL-4 (BioXCell). In Oxphos inhibition experiments, OXPHOSi (20nM) was added for the duration of polarization. After 72 hours, cells were stimulated with 1ng/mL PMA (Sigma), 1ug/mL Ionomycin (Invitrogen), and BrefeldinA 1000X (eBioscience) for 3 hours at 37C. Cells were stained with LIVE/DEAD™ Fixable Blue Dead Cell Stain Kit (Invitrogen) and/or monoclonal antibodies for flow cytometry (see Table 1). For intracellular staining, Cytofix/Cytoperm kit (BD Biosciences) was used for fixation and permeabilization. For detection of p-STAT3, cells were stimulated for 15 minutes with 1ng/mL PMA before fixation. Flow analysis was done on an Aurora (Cytek) and analyzed using FlowJo v.10.8.1 software.

For T cell co culture experiments, naïve T cells were polarized in 10mM glucose and Th2 polarizing conditions (detailed above) or 10mM galactose and Th17 polarizing conditions (detailed above) for 48h. Polarized Th2 or Th17 cells were co-cultured together (Th2/Th17) or with freshly isolated naïve T cells (Th2/naïve or Th17/naïve) in RPMI media (50:50 glucose and galactose), 20ug/mL IL-2, 10ng/mL PMA and either 5uM dexamethasone or vehicle control for 16h. Flow staining (see Table 1 for antibody information) and analyses were conducted as detailed above.

### Fibroblast-T cell co-culture

Fibroblasts were derived from WT mice via explant/transplant as previously described(76, 77). WT fibroblasts were plated at 50,000 cells per well in 8-well glass bottom chamber slides (Ibidi) and then incubated for 1 hour at 37. T cells were isolated as described above. After 1 hour, fibroblast media was removed and 1 million T cells in DMEM media (Gibco) containing either Th0 control or Th17 differentiating conditions (see above) were added to each well. For OPN-neutralizing conditions, anti-OPN (BioXCell) was used at 2ug/mL and was added to the media at time of plating. T cells were allowed to differentiate in co-culture with the fibroblasts for 72h. After 72h, co-cultures were fixed with 4% PFA prior to staining. Co-cultures were incubated in a blocking solution with 10% by volume skim milk powder, 10% donkey serum and Fc block for 1 hour at RT. For the fluorescent signal, Phalloidin-iFluor 647 (ab176759; Abcam) stain was completed at 1:1000 dilution for 45 minutes at RT. Nuclear stain is performed by using DAPI at 1:5000 dilution for 5 minutes at RT. Samples were preserved in ProLong Gold Antifade (Life Technologies) before imaging on a Nikon A1 inverted microscope using a 60X water objective.

### RNA sequencing

RNA was isolated with a Qiagen kit from HDM-induced IL4Rα^-/-^ or HDM-induced Balb/c mouse lungs or from flow-sorted populations, including T cells (CD45^+^ CD3^+^ CD44^+^CD62L^-^), eosinophils (CD45+ Ly6C+ SiglecF+), neutrophils (CD45+ CD11b+ GR1+ SigF-), macrophages (CD45+ CD11b+ Ly6C+ Mac3+ auto-fluorescence+) and stromal cells (CD45-) from HDM-induced IL4Rα^-/-^ or HDM-induced Balb/c animals. After Ilumina paired-end sequencing, reads were aligned to the GRCh38 reference genome. Low-complexity reads were eliminated from BAM files before conversion of SAM files and feature counts generation of read counts (with a minimum quality cutoff Phred > 10). DESeq2 was used to identify differentially expressed genes among the various sets of samples. P values were calculated by the Wald test and then adjusted for multiple test correction with the Benjamini-Hochberg algorithm. Genes were considered differentially expressed between two sample groups when the DESeq2 analysis resulted in an adjusted P-value of <0.05 with a fold change of 1.5. The sequencing data generated in this study is available on Gene Expression Omnibus *(to be deposited upon acceptance)*.

### qPCR

Total RNA was isolated using trizol (Invitrogen) and RNeasy Fibrous Tissue mini kit (Qiagen). Transcriptor First Stand cDNA kit (Roche) was used to prepare single stranded cDNA by reverse transcribing 5 µg total RNA. Samples were amplified using SYBR Green Supermix (Roche) and the primer pairs used are listed in Table 2. All samples were run in triplicate with mean values used for quantification and data shown are expression relative to L32 or GAPDH.

**Table 2:**
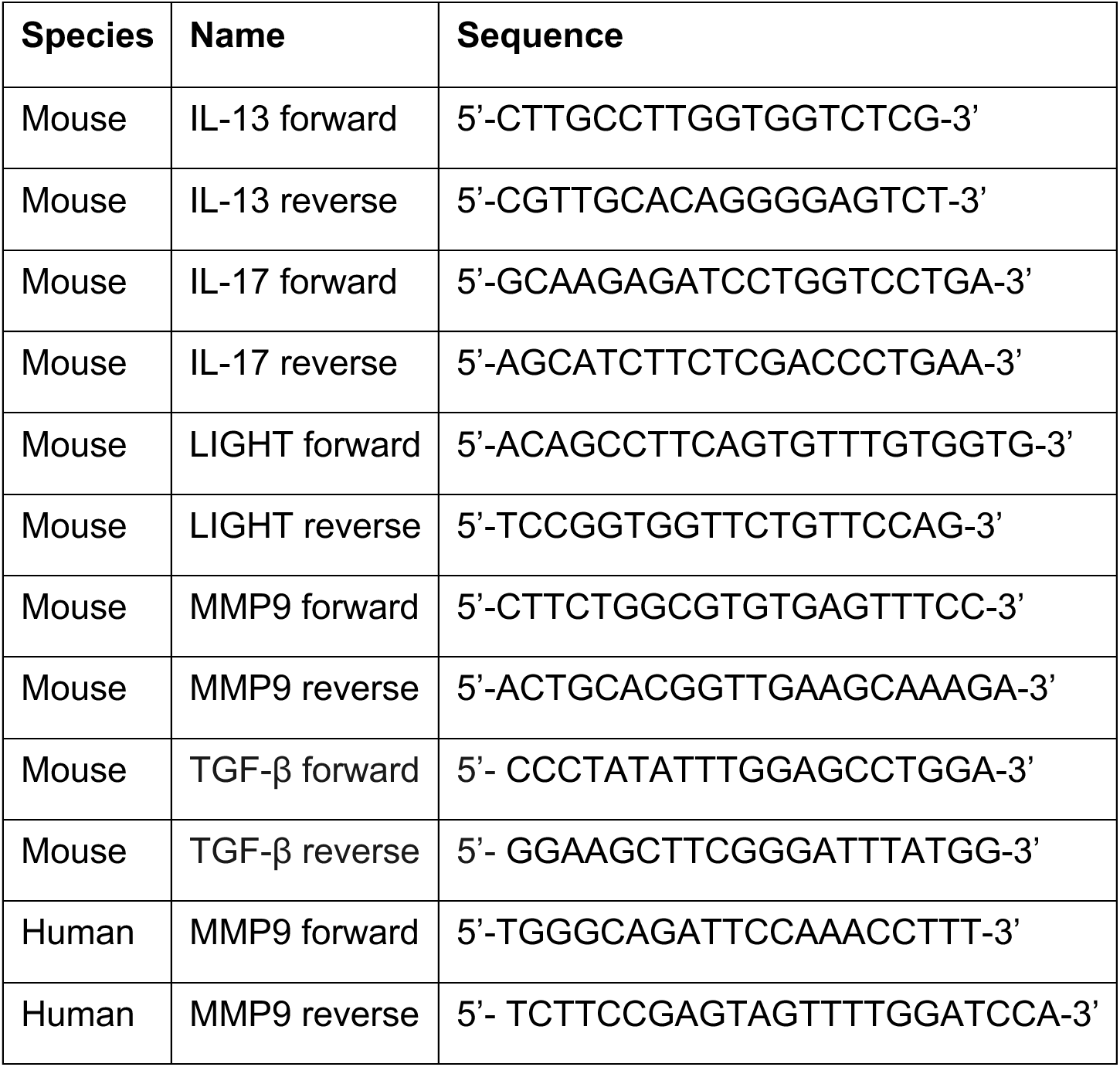
qPCR primers. A list of qPCR primer pair names and sequences used.

### Seahorse assay

Real-time OCR was measured using a XF-96 extracellular flux analyzer (Seahorse Bioscience). T cells from HDM-induced *Ilr4a^-/-^* or WT mice were sorted from the lung and washed twice with XF DMEM (pH 7.4) supplemented with 10mM glucose, 2mM L-glutamine, and 1mM pyruvate. Cells were seeded in the assay medium and attached in a Cell-Tak (Sigma) coated XF plate before resting for 1h at 37C prior to analysis. Mitochondrial stress test was performed following manufacturer instructions (Seahorse Bioscience) with final injection concentrations of 2uM Oligomycin, 1.5uM FCCP, 1uM Rotenone, and 1uM Antimycin A.

### Statistics

Statistical analysis was performed using GraphPad Prism v9.5.1 software. One way ANOVA or non-parametric Mann-Whitney U test was used where indicated. When One way ANOVA was used, multiple comparison was employed. A *P* value < 0.05 was considered statistically significant. Data is presented as means ± SEM. *\*P <* 0.05, *\*\*P <* 0.005, *\*\*\*P <* 0.0005, *\*\*\*\*P <* 0.00005.

## Acknowledgements

We thank Klaus Pfeffer and Stephanie Scheu for providing LIGHT^-/-^ mice. We thank Kacey Sachen, Rachel Soloff, Michael Croft, and Andrew McKnight for helping set up the model. We thank Paul Lembicz, Hailey Doerflein, and Therese Suchoski for assistance. We thank Katherine Baines, Megan Levings, Paritha Arumugam, and Georg Weber for supporting R.H. to get funding and sharing their expertise. We also thank the Bio-Imaging and Analysis Facility, Research Flow Cytometry Facility (including Mary Mullens for cell sorting), Integrated Pathology Research Facility (including Betsy DiPasquale and Lori Miller), and Veterinary Services Facility, as well as the Digestive Health Center (DHC) at CCHMC for their help. This work was supported by the US National Institutes of Health R01 AI177359 to R.H.

## Author Contributions

Conceptualization, R.H.; formal analysis, H.S. and R.H.; funding acquisition, R.H.; investigation, H.S., E.K., G.D., M.F., A.G., A.W. and R.H.; methodology, R.H., H.S.; project administration, R.H.; resources, E.K., M.F., H.C., M.K. and R.H.; supervision, R.H.; validation, H.S. and R.H.; visualization, H.S. and R.H.; writing– original draft, H.S. and R.H.; writing– review & editing, R.H., H.C. and M.K.

## Data and materials availability

RNAseq data presented in this study will be deposited on GEO as GSE dataset upon acceptance. This paper does not report original code. All other data are available in the main text of this paper.

